# Asymmetric localization of the cell division machinery during *Bacillus subtilis* sporulation

**DOI:** 10.1101/2020.07.22.216184

**Authors:** Kanika Khanna, Javier López-Garrido, Joseph Sugie, Kit Pogliano, Elizabeth Villa

## Abstract

The mechanistic details of bacterial cell division are poorly understood. The Gram-positive bacterium *Bacillus subtilis* can divide via two modes. During vegetative growth, the division septum is formed at the mid cell to produce two equal daughter cells. However, during sporulation, the division septum is formed closer to one pole to yield a smaller forespore and a larger mother cell. We use cryo-electron tomography to visualize the architectural differences in the organization of FtsAZ filaments, the major orchestrators of bacterial cell division during these conditions. We demonstrate that during vegetative growth, FtsAZ filaments are present uniformly around the leading edge of the invaginating septum but during sporulation, they are only present on the mother cell side. Our data show that the sporulation septum is thinner than the vegetative septum during constriction, and that this correlates with half as many FtsZ filaments tracking the division plane during sporulation as compared to vegetative growth. We further find that a sporulation-specific protein, SpoIIE, regulates divisome localization and septal thickness during sporulation. Our data provide first evidence of asymmetric localization of the cell division machinery, and not just septum formation, to produce different cell types with diverse fates in bacteria.

## Introduction

Bacterial cell division involves the invagination of the cellular membrane(s) and peptidoglycan (PG) cell wall to split the cell into two progeny cells. In most bacteria, cell division is initiated when the tubulin homologue and GTPase, FtsZ, polymerizes to form a ring-like structure, called the Z-ring, at the division site (1–3). The Z-ring serves as a scaffold onto which about a dozen other proteins assemble to form a mature cell division machinery called the ‘divisome’ (4). In the Gram-positive bacterium *Bacillus subtilis*, divisome assembly proceeds in two steps. First, proteins that interact directly with FtsZ and promote its association with the membrane are recruited to the divisome (5). One essential protein recruited at this stage is FtsA, an actin homologue which tethers FtsZ to the membrane via its conserved C-terminal amphipathic helix (6, 7). FtsA can also form protofilaments via polymerization in an ATP-dependent manner (8). Second, proteins involved in septal PG synthesis and remodeling are recruited to the divisome (5). Recent studies suggest that treadmilling by FtsZ filaments serves as a platform to drive the circumferential motion of septal PG synthases in both *B. subtilis* and *E. coli*, suggesting that this dynamic property of FtsZ filaments can regulate cell wall remodeling in the septal disc (9, 10).

*B. subtilis* is a unique model system to study cell division as it can divide via two modes. During vegetative growth, the septum is formed at the mid cell, producing two daughter cells of equal sizes (11). However, during sporulation, the division septum is formed closer to one pole to produce a smaller forespore and a larger mother cell. Subsequently, the forespore is engulfed by the mother cell, producing an internal endospore that, upon maturation, is released by mother cell lysis (12, 13). Vegetative and sporulating cells differ not only in the positioning of the division site but also in the amount of septal PG, with the medial vegetative septum being almost four times thicker than the polar sporulating septum (∼80 nm vs ∼23 nm) (14, 15). Another distinction is the presence of a membrane protein SpoIIE, which colocalizes with and directly interacts with FtsZ in only sporulating cells (16–18), but is absent in vegetative cells. Recent evidence suggests that SpoIIE localizes only on the forespore side of the polar septum at the onset of membrane constriction before being released into the forespore membrane upon septum formation (19–23). Certain *spoIIE* mutants also form thicker polar septa compared to wild type sporangia (14, 24). These findings posit that there may be differences in the divisome organization at the forespore and the mother cell side of the septum that play a role in regulating septal thickness. However, a mechanistic understanding of how cell division is regulated during *B. subtilis* sporulation remains elusive.

In this study, we used cryo-focused ion beam milling coupled with cryo-electron tomography (cryo-FIB-ET) to reveal the molecular architecture of the cell division machinery in *B. subtilis* during vegetative growth and sporulation. Our results demonstrate that FtsAZ filaments have distinct spatial organization during the two modes of growth and that SpoIIE regulates the positioning of FtsAZ filaments during sporulation. We further provide evidence that this distinct organization leads to a thinner polar septum during sporulation which has important physiological consequences during forespore engulfment. Our results pave the way for future studies in the field of bacterial cell division, giving rise to testable hypothesis on how bacteria regulate protein localization in space and time to carry out the critical process of cytokinesis under different gene expression programs.

## Results

### FtsAZ filaments in vegetative *B. subtilis*

Cryo-electron tomography (cryo-ET) is a method to obtain three-dimensional reconstructions of biological specimens. Previously, cryo-ET images of *E. coli* and *C. crescentus* revealed the presence of a series of dots at the division site corresponding to cross-sectional views of FtsZ filaments encircling the cell (6, 25). However, because of their increased thickness, *B. subtilis* cells were precluded from high-resolution tomography until recently, when we incorporated a cryo-FIB milling step in the workflow to provide mechanistic details about engulfment in *B. subtilis* sporangia (26, 27) (Fig. S1, see also Supplemental Information and Materials and Methods).

To get insights into the divisome architecture in *B. subtilis*, we first observed the leading edge of the invaginating septum in cryo-FIB-ET images of dividing vegetative cells (Fig. 1, S2). Our data revealed two series of dots at the division site that were distributed uniformly along the boundary of the leading edge: a membrane-proximal series of dots (∼6.5 nm from the invaginating membrane) and a membrane-distal series of dots (∼ 14 nm from the invaginating membrane) (Fig. 1C-E,H,L-N, S2A,B,D,E, Movie S1). Rotation of the 3D volume of the tomogram around the short axis of the cell demonstrated that each series of dots corresponded to filamentous structures of ∼ 3.5 nm diameter arranged next to each other, forming a bundle that spans the circumference of the cell (Fig. 1F-K, S2C,F, Movie S2). In the membrane-distal series of dots, we could easily resolve individual filaments spaced ∼ 5.5 nm apart while the filaments corresponding to the membrane-proximal series of dots were more diffuse (Fig. 1D,E). In a few data sets, we observed densities connecting the membrane-proximal and the membrane-distal bundles in a ladder-like arrangement (Fig. 1L-N) and additional densities connecting the membrane-proximal ring to the invaginating cellular membrane (Fig. S2F). In cryo-ET, contrast in images reflects variation in mass density across the biological specimen. Analysis of the intensities of the two rings revealed that the membrane-distal ring is denser and more continuous than the membrane-proximal ring which appears patchy (Fig. S2G).

**Fig. 1.**
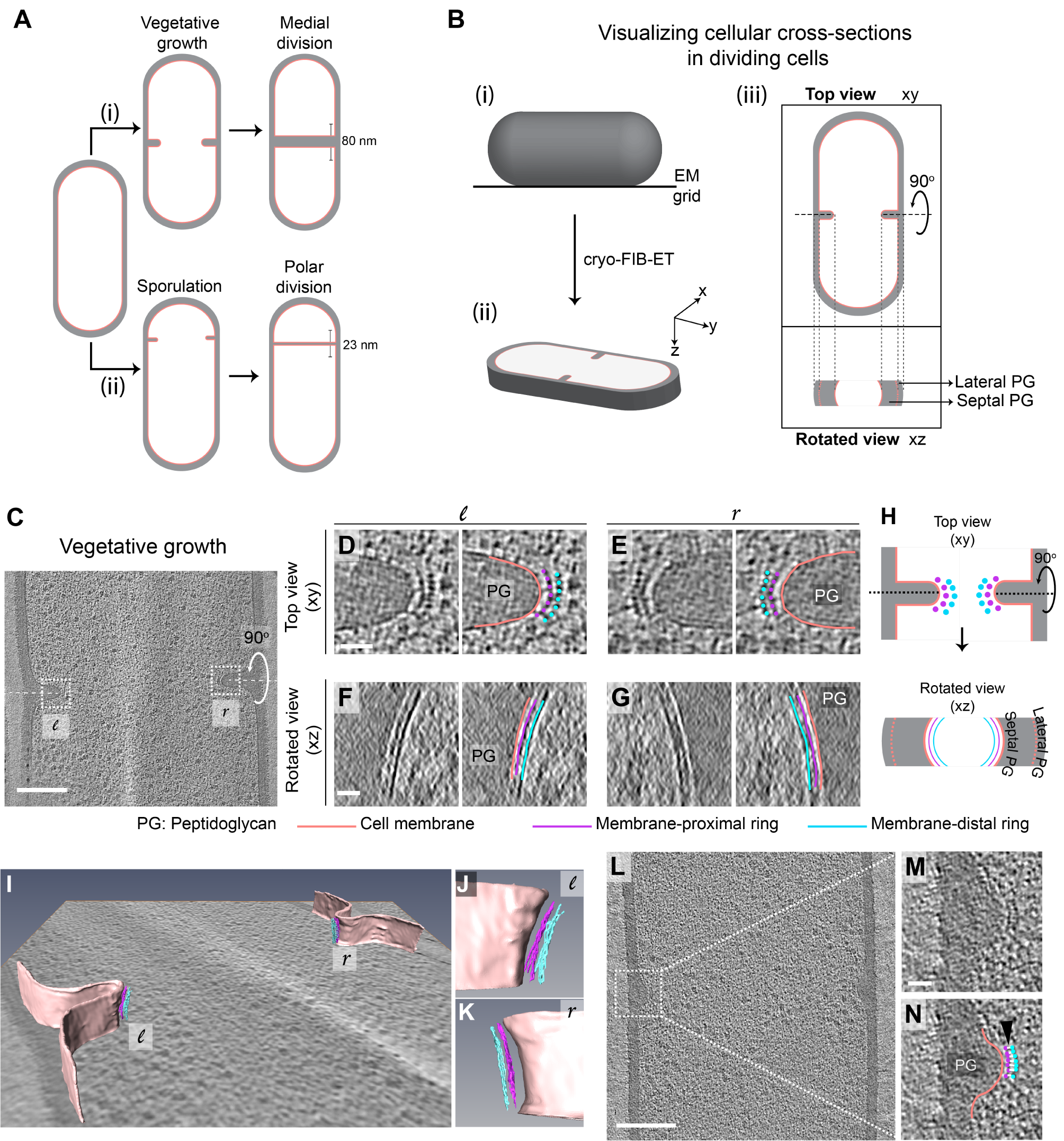
Architecture of divisome in vegetative *B. subtilis* cells. (A) Schematic of cell division in *B. subtilis* during (i) vegetative growth and (ii) sporulation. The thickness of septa upon their closure is indicated for both cases. (B) Schematic explaining visualization of cells in different planes in three-dimension (3D). (i) Initially, the rod-shaped *Bacillus* lies flat on an electron microscopy (EM) grid. (ii) Representation of 3D view of a cellular section obtained by cryo-FIB-ET in the xyz coordinate axis. x axis represents the length along the short axis of the cell, y axis represents the length along the long axis of the cell and z axis represents the height of the cellular specimen. (iii) Top panel, projection image of the cell in the xy coordinate plane (top view). Bottom panel, the corresponding projection image in the xz coordinate plane when the cell is rotated about its short axis by 90° (side/rotated view). The lateral and septal peptidoglycan (PG) are also indicated. (C) Slice through a tomogram of a dividing vegetative cell. The insets (*l* for left and *r* for right side of the septum) highlight the leading edges of the invaginating septum. (D) Left panel, zoomed-in view of the *‘l’* inset in (C) in the xy coordinate plane. Right panel, same as left with PG (grey), cell membrane (peach), membrane-proximal series of dots (pink) and membrane-distal series of dots (blue) highlighted. The same color scheme is followed throughout. (E) Left panel, zoomed-in view of the inset *‘r’* in (C) in the xy coordinate plane. Right panel, same as left with cellular parts highlighted. Left panel, view of the septal disc corresponding to (D) in the xz coordinate plane obtained by rotating the cell around its short axis by 90°. Right panel, same as left with PG, cell membrane, membrane-proximal ring and membrane-distal ring highlighted. (G) Left panel, view of the septal disc corresponding to (E) in the xz coordinate plane obtained by rotating the cell around its short axis by 90°. Right panel, same as left with cellular parts highlighted. (H) Schematic of the arrangement of the cytoskeletal machinery in dividing vegetative cells as seen in the xy and xz coordinate planes. All cellular parts detailed previously are highlighted in the same color scheme. (I) Annotation of the cell membrane and filaments corresponding to the membrane-proximal and the membrane-distal rings for the tomogram shown in (C). (J) and (K) represent zoomed-in views of the left (*l*) side and the right (*r*) side of the invaginating septum of the segmentation in (I) respectively. Scale bars are omitted from (I) – (K) owing to their perspective nature. (L) Slice through a tomogram of a dividing vegetative cell. The inset highlights the left side of the leading edge of the invaginating septum. (M) Zoomed-in view of the inset in (L) in the xy coordinate plane. (N) Same as (M), with cytoskeletal bundles and PG highlighted and ‘ladder-like’ connection between the membrane-proximal and the membrane-distal bands is shown by white lines. Scale bars for (C,L): 200 nm and for (D-G,M,N): 25 nm. See also Fig. S1, S2.

### FtsA and FtsZ comprise the membrane-proximal and the membrane-distal ring respectively

Our data suggest that the arrangement of the two cytoskeletal rings that mediate cell division in *B. subtilis* differs from Gram-negative bacteria like *E. coli* and *C. crescentus* where only a single bundle of FtsZ filaments is visible at a distance of ∼ 16 nm from the inner membrane (6, 25). In *E. coli*, another ring composed of FtsA filaments is visible at a distance of 8 nm from the membrane only when FtsA is overexpressed such that the ratio of FtsZ to FtsA alters from 5:1 in wild type to 1:1 in the modified strain (6). Based on these data and the knowledge that both FtsA and FtsZ form filaments in vivo and in vitro (3, 8), we hypothesized that in *B. subtilis*, the membrane-proximal ring is composed of FtsA filaments and the membrane-distal ring that of FtsZ filaments.

To unambiguously establish the identity of the two rings, we constructed a *B. subtilis* strain with an extra glutamine-rich linker region between the globular N-terminal domain of FtsZ and its C-terminal helix that binds FtsA (hereafter referred to as FtsZ-linker_Q-rich_, Fig. S3A-C). In a similar *E. coli* mutant strain, the Z-ring was further from the membrane by ∼ 5 nm compared to the wild type (6). In *B. subtilis* FtsZ-linker_Q-rich_, the distance of the membrane-proximal ring from the invaginating membrane remained unaltered compared to wild type, while the membrane-distal filaments were further from the membrane compared to wild type (17.5 ±1.1 nm vs 14± 0.6 nm) with a wide distribution ranging from ∼15 to ∼19 nm (Fig. 2A), establishing the membrane-distal filaments to be FtsZ.

**Fig. 2.**
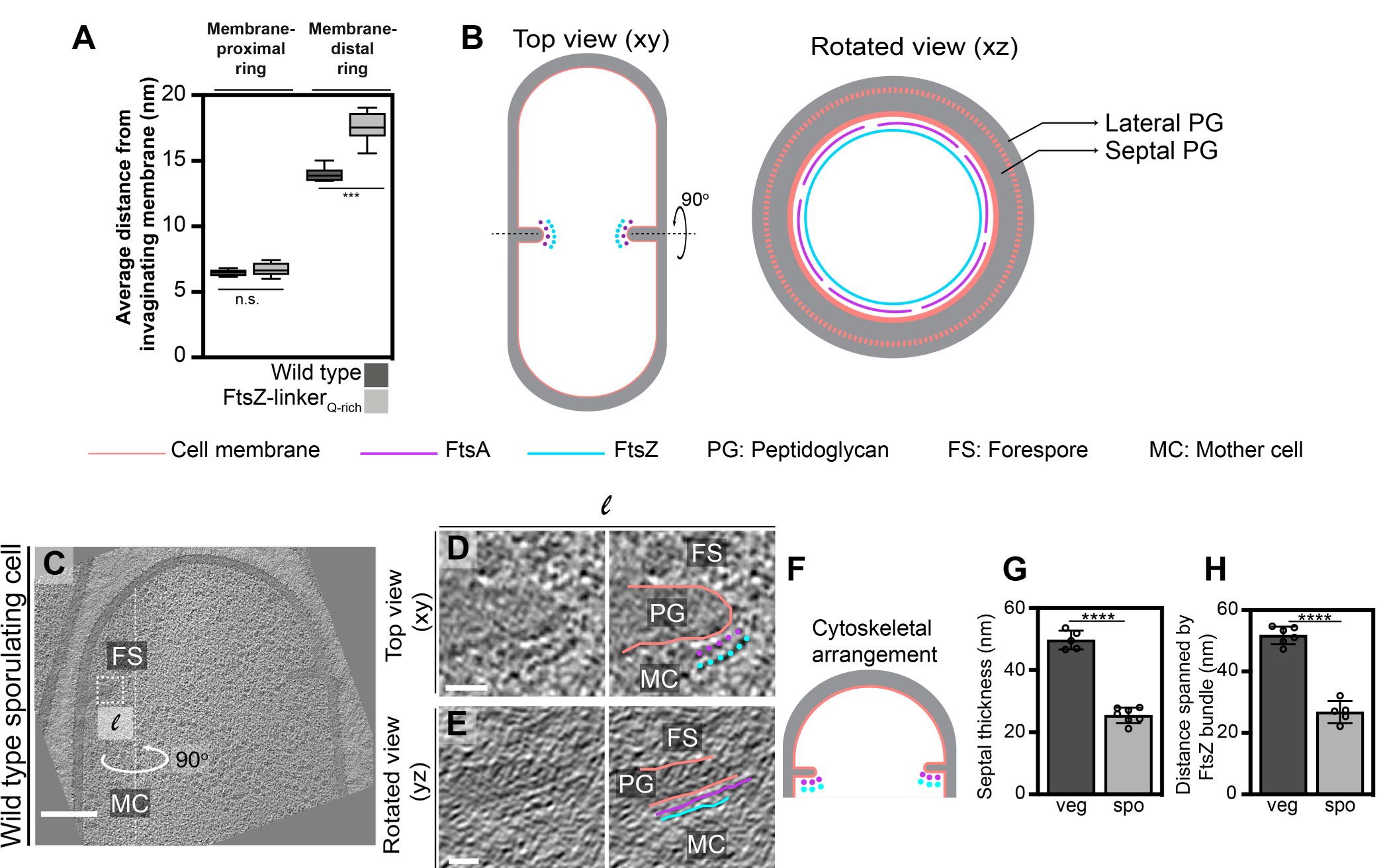
Identity of cytoskeletal filaments and their localization during sporulation. (A) Box-plot showing the distance of the membrane-proximal and the membrane-distal rings from the cell membrane in wild type and FtsZ-linker_Q-rich_ strains. Error bars indicate standard deviation (n.s.: p*>*0.05, ***: p *≤*0.001, unpaired t-test). (B) Schematic highlighting the arrangement and the identity of the cytoskeletal machinery with PG (grey), cell membrane (peach), FtsZ (blue) and FtsA (pink) highlighted. In the top view (xy coordinate plane), the cytoskeletal machinery is visible as two series of dots at the nascent septum. In the rotated view (xz coordinate plane), denser and more continuous Z-ring is tethered to the membrane via a patchy ring formed by FtsA filaments. (C) Slice through a tomogram of a dividing sporulating cell. (D) Left panel, zoomed-in view of the *‘l’* inset in (C) in the xy coordinate plane. Right panel, same as left with cell membrane (peach), FtsA bundle (pink) and FtsZ bundle (blue) highlighted. PG, forespore and mother cell compartments are also indicated. (E) Left panel, view of the septal disc corresponding to (D) in the yz coordinate plane obtained by rotating the cell around its long axis near the left side of the invaginating septum by 90°. Right panel, same as left with cellular parts and FtsAZ filaments highlighted. (F) Schematic of the arrangement of cytoskeletal machinery during sporulation. (G) and (H) Bar graphs depicting (G) septal thickness and (H) distance spanned by FtsZ bundle in wild type vegetative and sporulating cells. For both, error bars indicate standard deviation. (****: p*≤* 0.0001, unpaired t-test). Scale bar for (C): 200 nm and for (D,E): 25 nm. See also Fig. S3–S6.

In a few instances, we detected “stray” filaments at a distance of ∼12–20 nm from the membrane and away from the leading edge (black arrows, Fig. S3D-J). We speculate that such filaments correspond to FtsZ but without a corresponding FtsA filament to tether it to the membrane, as the in-corporation of the long linker likely affects the interaction between the two proteins (as also evidenced by longer filamented cells in the presence of the linker, Fig. S3A,B). In a few tomograms, we also observed doublets of individual filaments only in the membrane-distal band, an arrangement that was previously reported for FtsZ filaments in tomograms of dividing *E. coli* cells (6) (blue arrows, Fig. S3D,K-N), further demonstrating that only the membrane-distal ring corresponded to the Z-ring. In one tomogram, we observed an additional faint ring between the membrane-proximal and the membrane-distal rings (∼ 14 nm from the membrane) that might correspond to few FtsZ filaments that localize closer to the membrane likely due to the flexibility of the linker do-main (in yellow, Fig. S4A-C). We assigned the identity of the membrane-proximal ring to be FtsA filaments, as no other cell division proteins form filaments *in vivo* or *in vitro* and its distance to the membrane is comparable to that of FtsA filaments in *E. coli* upon overexpression (6). To further establish the identity of the filaments, we verified that the filament thickness, continuity along the septal plane, and distribution of intensity values of the rings in the FtsZ-linker_Q-rich_ mutant strain compared to the wild type were all similar (Fig. S4D). Thus, we conclude that the more continuous membrane-distal ring of FtsZ filaments is tethered to the membrane via a patchy membrane-proximal ring of FtsA filaments (Fig. 2B, Movie S3).

### FtsAZ filaments localize only on the mother cell side during sporulation

Next, we analyzed the divisome architecture in sporulating *B. subtilis* cells. During sporulation, the division site shifts closer to a pole as opposed to me-dial division during vegetative growth (Fig. 1A). We again observed two series of dots in dividing sporulating cells but remarkably, they were visible only on the mother cell side of the leading edge of the invaginating polar septum (Fig. 2C-F, S5, S6). Of note, the membranes at the leading edge of the nascent septum in sporulating cells were not as defined as in vegetative cells, suggesting they have a higher ratio of protein to membrane, as might be expected due to the presence of both transmembrane cell division proteins and sporulation-specific membrane proteins (Fig. 2C,D). In conclusion, our data suggest that FtsAZ filaments mediating cell division localize differently during vegetative growth and sporulation. During vegetative growth, they span the leading edge of the invaginating septum uniformly whereas during sporulation, they localize only on the mother cell side.

### FtsAZ filaments tracking the division plane may dictate septal thickness

Previous data demonstrated that upon closure, the polar sporulating septum is just one-fourth of the thickness of the medial vegetative septum (∼23 nm vs ∼80 nm) (14, 15). Recent studies suggest that FtsZ tread-milling drives the circumferential motion of septal PG synthases and condensation of the Z-ring increases recruitment of PG synthases in *B. subtilis* (9, 28). The observation that the sporulation septum is thinner than the vegetative septum and that FtsZ filaments localize differently during the two conditions led us to hypothesize that there is a direct correlation between the number of FtsZ filaments at the division site and the septal thickness during septal biogenesis.

To investigate this, we first measured the septal thickness in dividing sporulating and vegetative cells using cryo-FIB-ET images to determine if the sporulating septum is thinner than the vegetative septum as it is being formed or only upon closure as was previously reported (14, 15) (Fig. 2G). Our data showed that the thickness of the invaginating septa in vegetative cells was ∼50 ± 3 nm compared to ∼ 80 nm when the septum formation is completed, suggesting that during vegetative growth, either additional PG is incorporated into the septum upon its closure or the already present PG is remodeled and expands in volume after synthesis. On the other hand, the thickness of the invaginating septa in sporulating cells was almost half of that of vegetative cells (∼ 25 ±2.5 nm, Fig. 2G) during constriction. This suggests that even at the onset of cell division, the sporulating septum is thinner than the vegetative septum.

Next, we investigated if distinct localization of FtsZ filaments also translated into different number of FtsZ filaments tracking the division plane during vegetative growth and sporulation. It was often not possible to count the exact number of FtsZ filaments in cryo-FIB-ET data due to low signal-to-noise ratio at the division site, especially during sporulation given the molecular crowding at the leading edge of dividing septum. However, our data indicate that the distance between the individual FtsZ filaments is similar in both vegetative and sporulating septa, so we measured the distance spanned by the membrane-distal bundle of FtsZ dots as a proxy for the abundance of the cytoskeletal machinery during cell division. On average, FtsZ bundle spanned twice the length in vegetative cells compared to sporulating cells (∼ 51± 2.9 nm vs ∼ 27 ±3.6 nm, Fig. 2H), indicating that there are twice as many FtsZ filaments participating in cell division during vegetative growth as compared to sporulation. Based on these data, and previous results showing that the dynamic property of FtsZ filaments regulates the insertion of cell wall material during septal biogenesis (9, 28), we propose that the sporulation septum being thinner than the vegetative septum is likely a consequence of fewer FtsZ filaments tracking the division plane during sporulation.

### SpoIIE affects the localization of FtsAZ filaments during sporulation

We next probed factors leading to the distinct pattern of FtsAZ filament localization during vegetative growth and sporulation. We focused on dissecting the role of SpoIIE (or IIE), a sporulation-specific integral membrane protein proposed to regulate polar division for several reasons. First, IIE localizes to the dividing polar septum in an FtsZ-dependent manner (16–18). Biochemical evidence suggests that the central FtsZ-binding domain and possibly the N-terminal transmembrane domain of IIE play a role in its interaction with FtsZ (18, 19). Second, previous electron micrographs demonstrated that certain *spoIIE* mutants form thicker polar septa compared to wild type sporangia (14, 24). Third, recent evidence suggests that IIE preferentially localizes to the forespore side of the dividing septum, creating an asymmetry during polar division (20, 21, 23). Hence, we set out to determine if IIE regulates the divisome architecture during sporulation.

We first examined the divisome architecture in SpoIIE null mutant sporangia (*spoIIE*). Surprisingly, the divisome architecture in *spoIIE* is similar to that in vegetative cells, with two series of dots corresponding to FtsAZ filaments present uniformly along the leading edge in contrast to wild type sporangia where they were present only on the mother cell side (Fig. 3A-C,N). Also, at∼ 42± 4 nm, the invaginating polar septum in *spoIIE* sporangia was thicker than the wild type, implicating IIE in regulating both the localization of FtsAZ filaments and the septal thickness during sporulation (Fig. 3O). Of note, even though the length spanned by the bundle of FtsZ filaments in *spoIIE* was similar to that in wild type vegetative cells, *spoIIE* division septa were still ∼10 nm thinner, suggesting there may be other unidentified factors in addition to IIE that regulate the thickness of the polar septum (Fig. S7A).

**Fig. 3.**
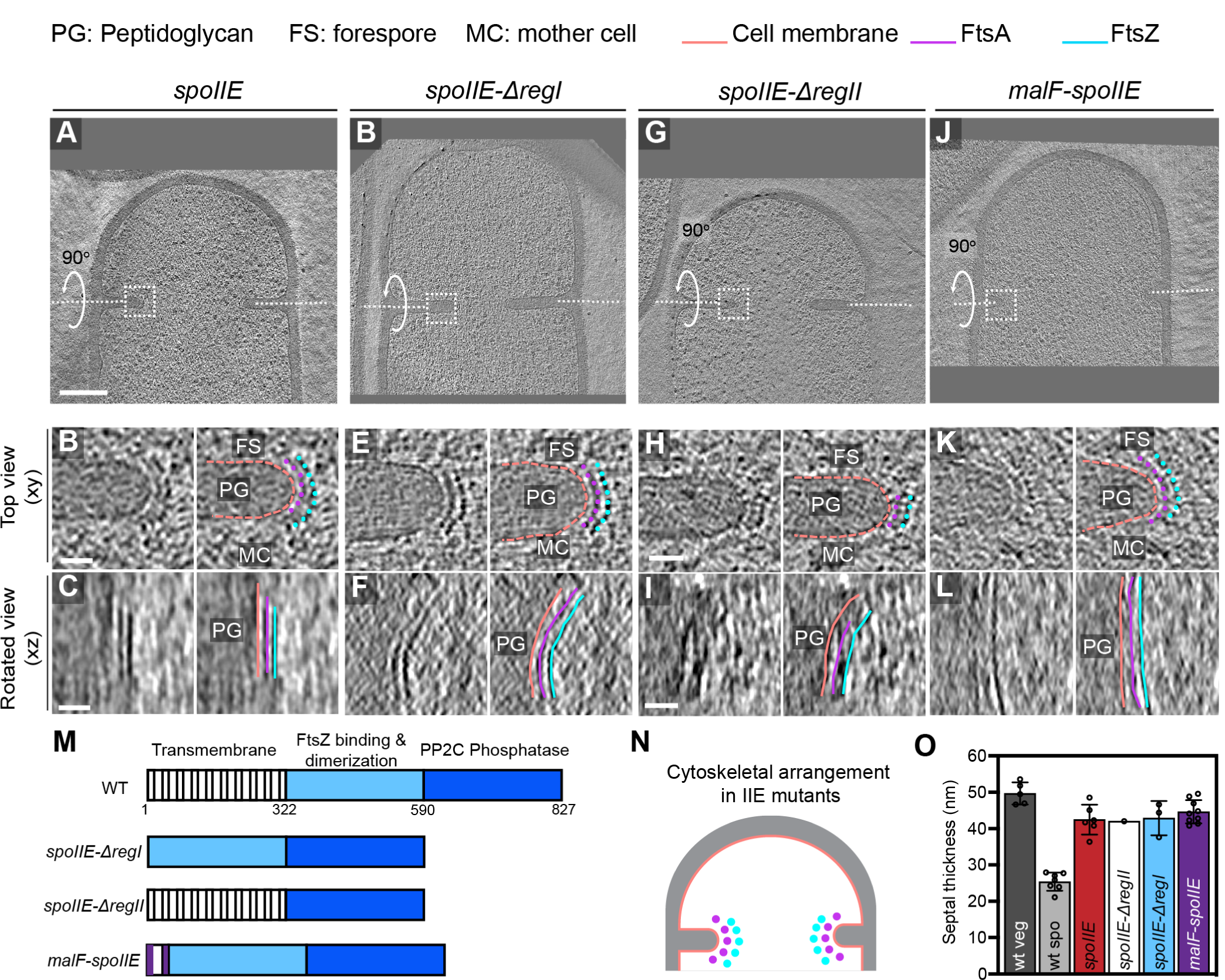
Localization of FtsAZ filaments in SpoIIE mutant sporangia. (A,D,G,J) Slice through a tomogram of a dividing (A) *spoIIE*, (D) *spoIIE-*Δ*regI*, (G) *spoIIE-*Δ*regII*, and (J) *malF-spoIIE* sporangia. Insets highlights the left side of the invaginating septum for each case. (B,E,H,K) Left panel, zoomed-in view of the insets in (A,D,G,J) respectively in the xy coordinate plane. Right panel, same as left with the peptidoglycan (PG), forespore (FS) and the mother cell (MC) compartments indicated in all. Cell membrane (peach), FtsA bundle (pink) and FtsZ bundle (blue) are also highlighted in all. (C,F,I,L) Left panel, view of the septal disc corresponding to (B,E,H,K) respectively in the xz coordinate plane obtained by rotating the cell around its short axis by 90°. Right panel, same as left with cellular parts and FtsAZ filaments highlighted. (M) Schematic highlighting different domains of SpoIIE and the mutants used in this study. (N) Schematic of the arrangement of the cytoskeletal machinery in SpoIIE mutant sporangia from (A)-(L). PG (grey), cell membrane (peach), FtsA bundle (pink) and FtsZ bundle (blue) are indicated. (O) Bar graph depicting the septal thickness in wild type vegetative and sporulating cells and SpoIIE mutant sporangia. Error bars indicate standard deviation. Each dot indicates a sample point. Scale bars for (A,D,G,J): 200 nm and for (B,C,E,F,H,I,K,L): 25 nm. See also Fig. S7.

IIE has three domains: an N-terminal transmembrane domain with 10 membrane-spanning segments (region I), a central domain that promotes oligomerization of IIE and its interaction with FtsZ (region II) and a C-terminal phosphatase domain that is involved in the activation of the forespore-specific transcription factor σ^*F*^ after septum formation (region III) (19). Since regions I and II have been implicated in regulating the interaction of IIE with FtsZ (18, 19), we acquired cryo-FIB-ET images of the following IIE mutant strains: (1) deletion of region I (or *spoIIE-*Δ*regI*), (2) deletion of region II (or *spoIIE-*Δ*regII*), and (3) replacement of 10 membrane-spanning segments of region I by 2 membrane-spanning segments from *E. coli* MalF (or *malF-spoIIE*). These strains also had a deletion in the *spoIIA* operon to uncouple the role of IIE in polar division from σ^*F*^ activation. Our cryo-FIB-ET data demonstrated that the organization of FtsAZ filaments at the polar septum and the septal thickness in all the three mutant strains is similar to *spoIIE* sporangia (Fig. 3D-N). These data indicate that both the transmembrane and the FtsZ-binding domain are essential to regulate the localization of FtsAZ filaments and septal thickness during sporulation.

### IIE affects localization of FtsA on the forespore side of the septum

IIE is present exclusively on the forespore side of the septum even before the onset of membrane constriction (20, 21, 23) and our data show that both the membrane-spanning and the FtsZ-binding domains of IIE mediate differential localization of FtsAZ filaments during sporulation. Since IIE directly binds FtsZ, one possible way IIE could affect the localization of FtsZ filaments on the forespore side would be by competing directly with other cell division proteins that bind to FtsZ, including FtsA. Hence, we hypothesized that the interaction of IIE with FtsZ may disrupt the interaction of FtsA with FtsZ on the forespore side. This could either be a result of direct competition faced by FtsA from IIE for the binding site on FtsZ, or due to possible membrane curving on the forespore side mediated by IIE oligomers upon insertion of their large membrane-spanning segment as has been previously reported in protein-packed environments (29). In support of this, using time-lapse microscopy and structured-illumination microscopy, in a few cells we observed two puncta of fluorescently tagged FtsA at the division septum, one that constricts with the invaginating membrane and another that remains at the intersection of the lateral and the septal cell wall at the forespore edge (Fig. 4A-C). These data suggest that FtsA is present but likely not incorporated into the cell division machinery on the forespore side of the septum. This, in turn, may affect the assembly of the complete divisome on the forespore side of the invaginating septum so that cytokinesis exclusively proceeds from the mother cell side (Fig. 4D). We anticipate that future biochemical and genetic studies based on these data may conclusively establish the mechanism by which IIE interacts with different components of the divisome to regulate cell division during sporulation.

**Fig. 4.**
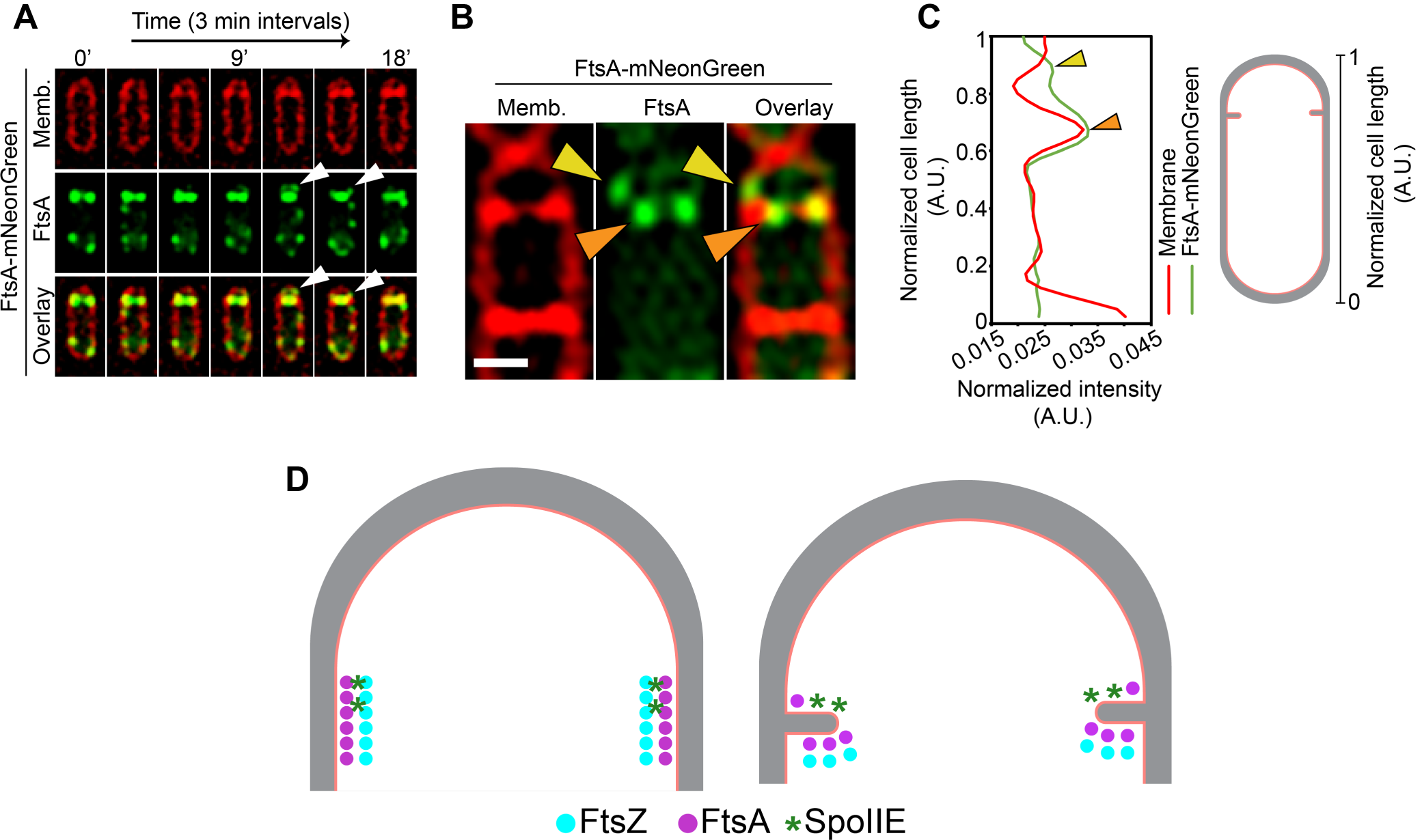
Role of SpoIIE in modulating FtsAZ filament localization. (A) Time-lapse fluorescent microscopy of a dividing sporangia with FtsA in green and membranes in red. Images are taken every 3 minutes. Between 12 and 15 minutes, two FtsA rings are visible, one that constricts and the other that seemingly stays behind at the forespore side of the septum. White arrows indicate FtsA on the forespore side that does not constrict. (B) Structured-illumination microscopy of a dividing sporangia with FtsA in green and membranes in red. The two puncta corresponding to FtsA are more clearly visible, one that constricts (orange) and the other that remains on the forespore side (yellow arrow). (C) Line graph showing the normalized intensity of the membrane (red) and FtsA-mNeonGreen (green) signal along the normalized length of the cell. Two peaks corresponding to two puncta of FtsA in (B) are indicated by orange and yellow arrows respectively. (D) A possible model of how SpoIIE (green asterisks) affects the localization of FtsA (pink) and FtsZ (blue) filaments during sporulation (top view of the cell shown). PG (grey) and cell membrane (peach) are also highlighted. IIE molecules are preferentially present on the forespore side and hence bind to FtsZ on the forespore side (brown asterisks). This may prevent binding of FtsA filaments (and likely other cell division proteins) to FtsZ on the forespore side and they stay behind at the forespore edge, while FtsAZ filaments on the mother cell side constrict. Scale bar for (A,B): 1 *µ*m.

## Discussion

In this study, we elucidate the differences in the architecture of the cell division machinery in *B. subtilis* during vegetative growth and sporulation and how they impact septal PG thickness. First, we show that during vegetative growth, two cytoskeletal rings localize uniformly around the leading edge of the invaginating medial septum to orchestrate cell division, a membrane-proximal ring of FtsA filaments that tethers the membrane-distal ring of FtsZ filaments to the membrane (Fig. 1, 2A,B). Second, we demonstrate that during sporulation, FtsAZ filaments are present only on the mother cell side of the invaginating polar septum (Fig. 2C-F). Third, our data suggest that during septal biogenesis, the number of FtsAZ filaments tracking the sporulating septum is approximately half of that in the vegetative septum which likely gives rise to a thinner sporulation septum by mediating the circumferential motion of fewer PG synthases around the division plane during sporulation (Fig. 2G,H). Finally, we show that a sporulation-specific protein SpoIIE is essential to regulate the localization of FtsAZ filaments and septal thickness during sporulation (Fig. 3, 4).

Most of our understanding of bacterial cytokinesis comes from studies in model Gram-positive and Gram-negative bacteria, namely *Bacillus subtilis* and *Escherichia coli*, respectively. Although the overall mechanism of cytokinesis is conserved in that a Z-ring that recruits other components of the divisome is formed at the division site, there are subtle differences with regards to the spatial and temporal regulation of the process and the composition of the divisome for both bacterial species. Previously, only a single cytoskeleton ring of FtsZ was observed using cryo-ET in Gram-negative bacteria including *E. coli* and *C. crescentus* (6, 25). However, we observed two rings corresponding to FtsA and FtsZ filaments in the Gram-positive bacterium *B. subtilis* during both vegetative growth and sporulation, even though the ratio of FtsZ to FtsA is same (5:1) in *B. subtilis* and *E. coli* (30, 31) and there are enough molecules of FtsA to form a complete circumferential ring in both. But an A-ring is apparent only in *B. subtilis*, although it appears patchy compared to the mostly continuous Z-ring. One possible explanation for this phenomenon could be attributed to the fact that cell division in *B. subtilis* involves constriction of a septal disc composed of thicker septal PG (∼ 27 nm during sporulation and ∼50 nm during vegetative growth). On the other hand, cell division in *E. coli* involves constriction of a septal disc composed of only a single layer of ∼4 nm thick PG. Hence, cytokinesis in *B. subtilis* will likely depend on scaffolding action of multiple FtsAZ filaments to drive the circumferential motion of more PG synthases. The lateral spacing between FtsZ filaments is ∼5.5 nm in *B. subtilis* which is slightly lower than the values reported in *E. coli* and C. crescentus (∼6.5-8 nm) (6). Also, the distances of the Z-ring and the A-ring from the membrane in *B. subtilis* (∼14 nm and ∼6.5 nm respectively) are slightly shorter than the corresponding values for *E. coli* (∼16 nm and∼ ∼ 8 nm respectively) (6). It is possible that a tighter bundling of FtsAZ filaments with each other and with the membrane in *B. subtilis* aids in efficient remodeling of the thicker septal PG. Whether these mechanisms of Z-ring and A-ring formation are also conserved in other sporulating and non-sporulating Gram-positive bacteria is yet to be explored.

Our cryo-FIB-ET data indicate that the Z-ring appears mostly continuous in the cellular sections we imaged, probably consisting of shorter overlapping filaments. 9 out of 16 tomograms of dividing vegetative cells that we captured showed more or less continuous Z-rings. The other tomograms were of significantly lower quality as they were either too thick or their orientation with respect to the tilt axis did not yield high-resolution information about the Z-ring. A previous cryo-ET study also revealed that the Z-rings are continuous in both *E. coli* and C. crescentus (6). However, superresolution microscopy data recently suggested that the organization of the Z-ring in both *B. subtilis* and *E. coli* is patchy with regions of low fluorescence intensity that indicate gaps in the ring (32, 33). This is in contrast to cryo-ET data which provides information at a resolution of a few nanometers. So far, it is difficult to ascertain the lengths of individual overlapping filaments unambiguously from our data and it is indeed possible that we do not see the gaps in our data as they may be present in the remainder of the cellular volume that has been ablated by cryo-FIB milling or not sampled due to missing wedge issue associated with data collection (Fig. S1). However, it is unlikely as the ring appears continuous in multiple tomograms, all of which capture different parts of the septal disc.

Our data demonstrate that during sporulation, not only is there asymmetry associated with the positioning of the division septum but also with the positioning of the division machinery as FtsAZ filaments are only present on the mother cell side of the invaginating septum. In fact, even the number of filaments tracking the division plane are half in sporulation compared to vegetative growth and so is the septal thickness. A thinner septum could be physiologically relevant to sporulation in two possible ways. First, many protein complexes are proposed to form channels across both the septal membranes during sporulation (34, 35). A thinner septum can facilitate interaction between the channel components in the forespore and the mother cell more efficiently. Second, a thinner septum may be easier to bend to facilitate migration of the mother cell membrane during engulfment.

Asymmetric cell division is a common developmental theme in eukaryotes to generate progeny with diverse cellular fates (36). During asymmetric cell division in *Caenorhabditis elegans*, the positioning of the sperm upon its entry in a newly formed zygote defines the posterior cell pole. A sperm associated signal then initiates the polarized distribution of PAR proteins which also regulate the alignment of the spindle by displacing it towards the posterior axis to ensure proper segregation of cell fate determinants. The asymmetric cleavage then produces two cells of different sizes and fates, the larger anterior cell and the smaller posterior cell (36). Asymmetric division is also important for nervous system development in *Drosophila melanogaster* wherein neuroblasts, the precursors of the central nervous system divide asymmetrically to produce larger apical daughter which remains a neuroblast and a smaller basal ganglion mother cell (GMC). Polarity is established by differential distribution of cell fate determinants to the basal and apical cortex. In this case, the spindle itself becomes asymmetrical with apical microtubules elongating and basal microtubules shortening and the midbody positions itself asymmetrically between the spindle poles (36).

In these and other cases of asymmetric cell division, the general theme is the establishment of cell division machinery along an axis of polarity and distribution of cell fate determinants in a polarized manner along this axis. In case of *B. subtilis* sporulation, an axis of polarity is likely established when the division plane shifts from medial to polar position and cell fate determinants are differentially distributed between the future forespore and the mother cell compartments at the onset of membrane constriction. In this study, we have identified SpoIIE as a cell fate determinant that restricts FtsAZ filaments on the mother cell side for polar division by preferentially localizing on the forespore side of the invaginating septum. From our data, we hypothesize that there are other, as of yet unidentified cell fate determinants that either recruit IIE or are recruited by IIE to the septum. This would explain why in the absence of IIE, even though the same number of FtsZ filaments track the septum as in the wild type vegetative septum, the invaginating septum is still ∼10 nm thinner (Fig. S7A).

Our data provide a significant advancement in the understanding of the organization of the divisome, its regulators and its role in septal PG synthesis in *Bacillus subtilis*. Future efforts to identify the molecular arrangement of FtsZ and its regulators inside the cell such as IIE, will aid in development of new antimicrobials targeting the cell division machinery in important pathogens.

## ACKNOWLEDGEMENTS

This work was supported by the National Institutes of Health R01-GM057045 (to K.P. and E.V.), and the National Science Foundation MRI grant NSF DBI 1920374 (to E.V.). We acknowledge the use of the UC San Diego Cryo-EM facility, which was built and equipped with funds from UC San Diego and an initial gift from Agouron Institute, and of the San Diego Nanotechnology Infrastructure (SDNI) of UC San Diego, a member of the National Nanotechnology Coordinated Infrastructure, supported by the NSF grant ECCS-1542148. We thank Dr. Marcella Erb for help with 3D-SIM experiments, and members of the Pogliano and Villa labs for their many insightful comments. We thank Richard Losick, Kumaran Ramamurthi and Ethan Garner for the gift of strains. We thank the Henriques Lab for the bioRxiv LATEX template.

## AUTHOR CONTRIBUTIONS

K.K., J.L.G, K.P. and E.V. designed the research. K.K. conducted all the experiments. All authors discussed the results and contributed to data analysis. K.K. wrote the paper with input from all other authors.

## Materials and Methods

### Strains and culture conditions

*Bacillus subtilis* PY79 background was used for all strain constructions. A list of strains used in the study is provided in Table S1. All the strains were routinely grown in LB plates at 30°C overnight.

For sample preparation, cells were either grown in LB media at 30°C to OD600∼ 0.5 for vegetative growth or by growing in 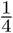 diluted LB to OD600 ∼0.5-0.8 and then resuspending in A+B media at 37°C for inducing sporulation. For wild type sporangia, samples were prepared at an early enough time point of ∼ T1.5-1.75 hours after inducing sporulation to capture a mixed population of dividing cells undergoing the formation of polar septum (sporulating cells) or medial septum (vegetative cells). For other mutant sporangia, samples were also collected at ∼T1.5-1.75 hours after inducing sporulation.

### Cryo-FIB-ET workflow

QUANTIFOIL R2/1 200 mesh holey carbon Cu grids (Quantifoil Micro Tools) were glow discharged using PELCO easiGlow (Ted Pella). 7-8 *µ*l of diluted liquid culture was deposited onto the grids which were mounted to a manual plunger (Max Planck Institute of Biochemistry) and manually blotted using Whatman No. 1 filter paper for 3-4 seconds from the side opposite to where the cells were deposited such that the cells formed a homogeneous monolayer on the grid surface. The grids were then plunge-frozen in a 50-50 mixture of liquid ethane and propane (Airgas) cooled to liquid nitrogen temperature and subsequently clipped onto Cryo-FIB Autogrids (Thermo Fisher Scientific). All subsequent transfers were performed in liquid nitrogen.

Frozen-hydrated cells were micromachined either inside Scios or Aquilos DualBeam instruments each equipped with a cryo-stage (Thermo Fisher Scientific). An integrated gas injection system (GIS) was used to deposit an organometallic platinum layer inside the FIB chamber to protect the specimen surface and avoid uneven thinning of cells. In case of specimens prepared in Aquilos, the specimen was also sputter coated with inorganic platinum layer prior to GIS deposition to prevent charging during imaging. FIB milling was then performed using two rectangular milling patterns to ablate the top and the bottom parts of the cells in steps of decreasing ion beam currents at a nominal tilt of 11°-15°which translates into a milling angle of 4°-8°. Initial rough milling was performed using ion beam currents of 0.5-0.3 nA followed by intermediate milling at 0.1 nA or 50 pA. Finally, fine milling was conducted at 30 pA or 10 pA. In each of the steps, the distance between the two rectangular milling patterns was sequentially decreased to get ∼ 100-300 nm thick lamellas.

Tilt series were collected using SerialEM software (37) either in 300-keV Tecnai G2 Polara or in 300-keV Titan Krios (Thermo Fisher Scientific), both equipped with a Quantum post-column energy filter (Gatan) and a K2 Summit 4k x 4k pixel direct detector camera (Gatan). The TEM magnification corresponded to a camera pixel size of either 0.612 nm (for data acquired on Polara) or 0.534 nm or 0.426 nm (for data acquired on Krios). The tilt series were usually collected from −61°to + 61°depending on the quality of the specimen with an increment ranging from 2°-3°and a defocus of −5 *µ*m following either the bidirectional or the dose-symmetric tilt scheme in low dose mode. For both schemes, the zero of the tilt series was defined by taking into account the pre-tilt of the lamella. The K2 detector was operated in counting mode and images divided into frames of 0.1s. The cumulative dose for each tilt series was ∼50-150 e^*−*^/Å^2^.

### Tomogram reconstruction and segmentation

Motion-Cor2 (38) was used to align the frames and dose-weigh the tilt series according to the cumulative dose. Subsequent alignment of the tilt series was done in IMOD (39) using patch-tracking in the absence of fiducials and the tomograms reconstructed using weighted back-projection method. For purposes of representation and segmentation, tomograms were binned 3x or 4x. Semiautomatic segmentation of the membranes was performed using TomosegmemTV (40) followed by manual refinement in Amira software package (Thermo Fisher Scientific). FtsAZ filaments were manually traced in Amira.

### Distribution of intensities of cytoskeletal rings

For Fig. S2G, S4D, the membrane-proximal and membrane-distal cytoskeletal rings were masked using Adobe Photoshop and the corresponding intensity values for each pixel were extracted using a custom-built MATLAB script. The distribution of these values for the two rings were depicted as box-and-whisker plot alongside the values corresponding to cell membrane, cytoplasm and cell wall (PG) that were used as controls.

### Distances between cytoskeletal rings and membrane

For Fig. 2A, a medial slice corresponding to the orthogonal view (xz) of the respective tomograms was taken from the z-stack wherein the cytoskeletal rings were clearly visible. The distances of the ring from the cell membrane were then calculated using the Fiji plugin ‘points to distance’.

### Fluorescence microscopy for batch cultures

12 *µ*l of sample were taken at indicated time points and transferred to 1.2% agarose pads that were prepared using either sporulation resuspension media (for sporulating cultures) or 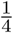 LB media (for vegetative growth). Membranes were stained with 0.5 *µ*g/ml of FM4-64 (Thermo Fisher Scientific) that was added directly to the pads. An Applied Precision DV Elite optical sectioning microscope equipped with a Photometrics CoolSNAP-HQ2 camera was used to visualize the cells. The images were deconvolved using SoftWoRx v5.5.1 (Applied Precision). For all fluorescence images, the medial focal plane of the image is shown. Excitation/ emission filters were TRITC/CY5 for membrane imaging and FITC/FITC to visualize GFP or mNeonGreen signal.

### 3D-Structured Illumination Microscopy (3D-SIM)

For Fig. 4B, cells were grown as indicated above and stained with 0.5 *µ*g/mL FM4-64. An Applied Precision/GE OMX V2.2 Microscope was then used to image them. Raw data were sequentially taken by SI-super-resolution light path to collect 1.5 mm thick specimens in 125 nm increments in the z-axis with compatible immersion oils (Applied Precision). Standard OMX SI reconstruction parameters were then used in DeltaVision SoftWoRx Image Analysis Program to reconstruct the images. To plot the graph in Fig. 4C, the data corresponding to membrane and mNeonGreen intensity was grouped into smaller bins of approximately equal area along the normalized length of the cell (as indicated in the inset) using a custom-built MATLAB script.

### Spore titer assay

Strains were grown in triplicates in 2 ml of DSM media for 24 hr at 37°C followed by heating at 80°C for 20 min. Then, serial dilutions for each strain were prepared and spotted on LB plates. Number of colonies were then used as a marker to calculate spore titers.

### Calculating GFP intensity

For Fig. S7C, ∼15-16 cells with clear evidence of septal biogenesis were manually selected for each strain and all the images were aligned such that the polar septum lies in the same orientation for all cells as shown in the inset of Fig. S7C(i). To get the linear profile of GFP intensity for each cell, the data was grouped into smaller bins of approximately equal areas using a custom-built MATLAB script. Normalized GFP intensities were then plotted for each cell along its normalized length as depicted by different curves in the graphs.

## Supplemental Information

**Table S1.**
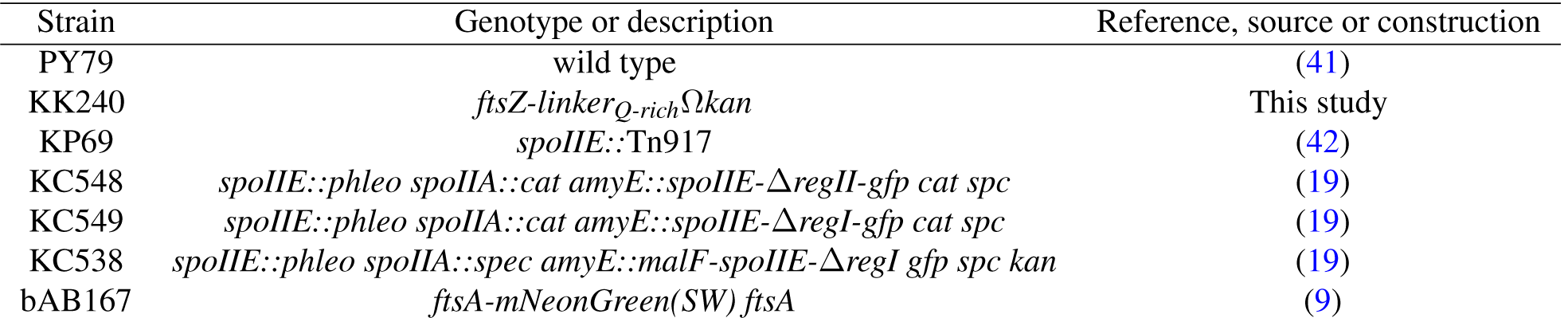
Strains used in this study.

### Construction of KK240

Constructed by transformation of pKK238 (FtsZ-linker_Q-rich_Ωkan) into PY79. pKK238 was constructed by removing ssrA tag of pJLG142 and incorporating Q-rich linker from FtsN of *E. coli* at the C-terminus of FtsZ. FtsZ-linker_Q-rich_ sequence: (Q-rich linker region highlighted in yellow and red)

MLEFETNIDGLASIKVIGVGGGGNNAVNRMIENEVQGVEYIAVNTDAQALNLSKAEVKMQIGAKLTRGLGAGANPEVGKKAAEESKEQIEEALKGADMVFVTAGMGGGTGTGAAPVIAQIAKDLGALTVGVVTRPFTFEGRKRQLQAAGGISAMKEAVDTLIVIPNDRILEIVDKNTPMLEAFREADNVLRQGVQGISDLIATPGLINLDFADVKTIMSNKGSALMGIGIATGENRAAEAAKKAISSPLLEAAIDGAQGVLMNITGGTNLSLYEVQEAADIVASASDQDVNMIFGSVINENLKDEIVVTVIATGFIEQEKDVTKPQRPSLNQSIKTHNQSVPKREPKREEPQQQNTVSRHTSQPARQQPTQLVEVPWNEQTPEQRQQTLQRQRQAQQLAEQQRLAQQSRTTEQSWQQQTRT-SQAAPVQAQPRQSKPASSQQPYQDLLQTPAHTTAQSKPQQD

**Fig. S1.**
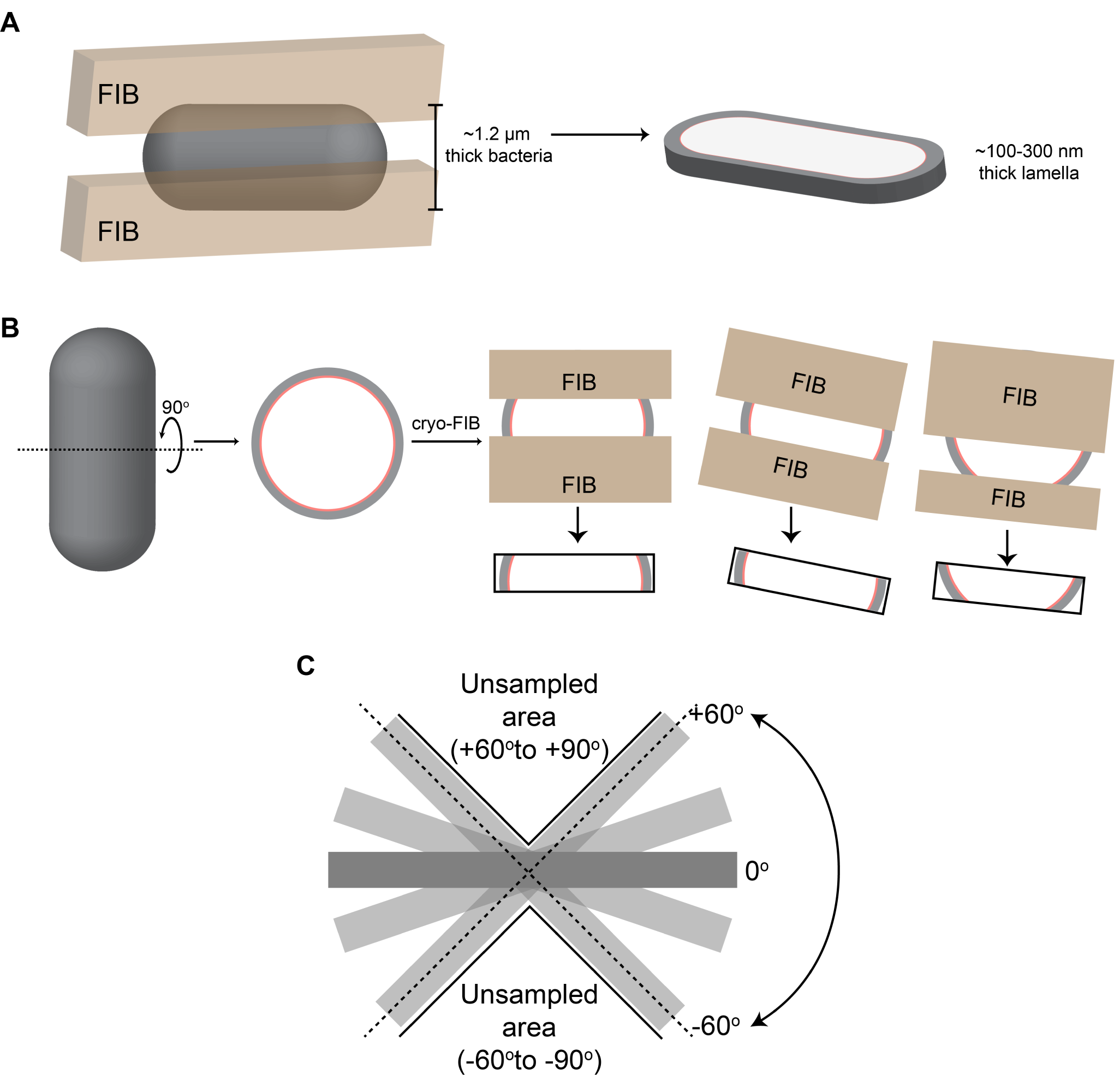
Cryo-FIB milling workflow and analysis. (A) Schematic of a rod-shaped cell (grey) subjected to cryo-focused ion beam (FIB) milling. Two parallel beam of gallium ions (brown) ablate the cellular material from the top and the bottom leaving a thin slice (∼100-300 nm thick) for imaging using cryo-electron tomography (cryo-ET). (B) Schematic depicting different sections of the septal disc that can be captured when a rod-shaped cell is FIB-milled depending on the milling angle, orientation of the cell on the EM grid and orientation of the cell with respect to the tilt axis. (C) Schematic depicting the missing wedge issue in cryo-ET workflow. Since we are only able to image from ∼ ±60°, the cellular volume from −60° to −90° and from +60° to +90° remains unsampled leading to missing information for these areas.

**Fig. S2.**
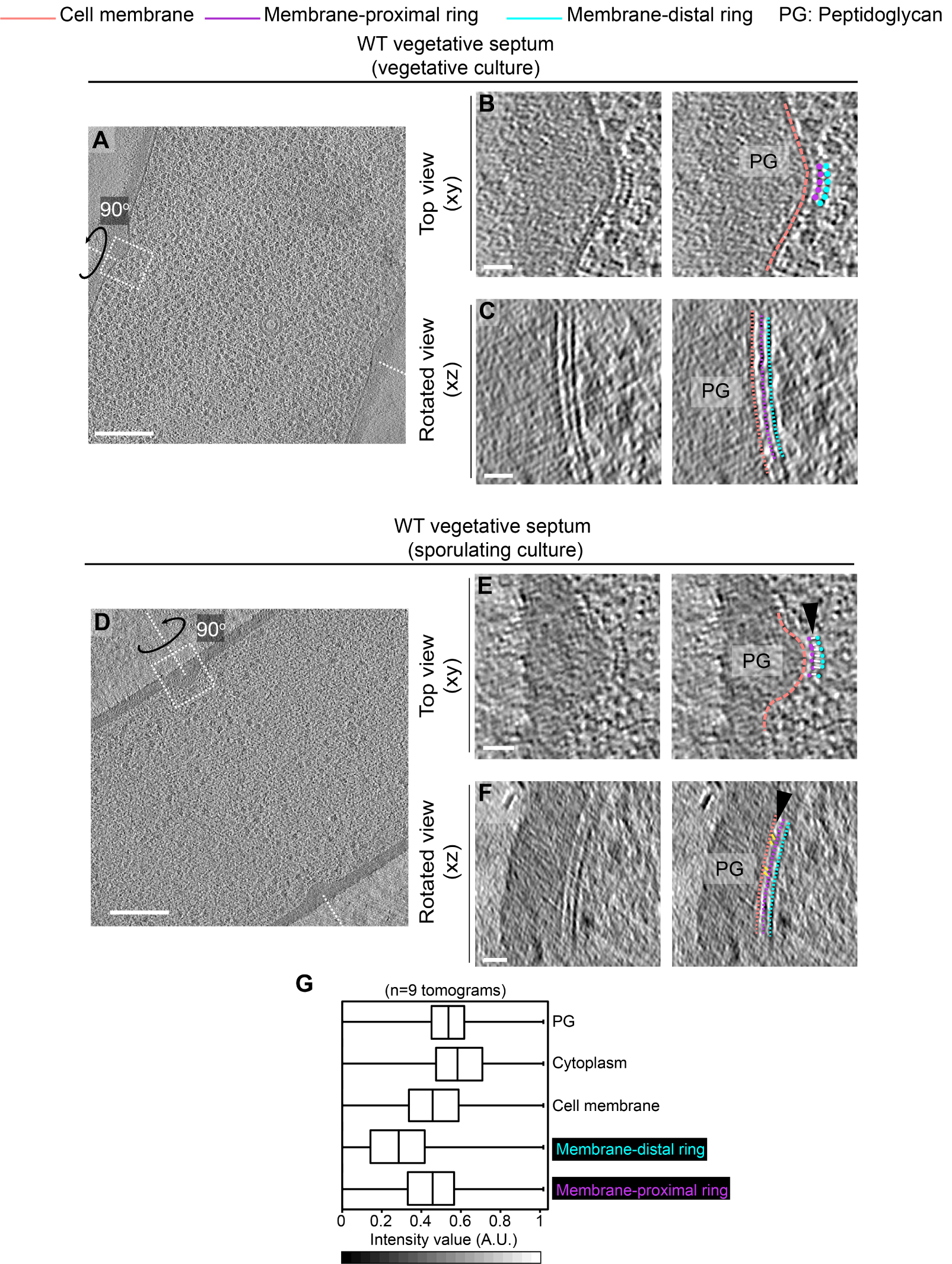
Cytoskeletal filaments in vegetative cells. (A) Slice through a tomogram of a dividing vegetative cell under vegetative culture conditions (see Materials and Methods). The inset highlights the left side of the leading edge of the invaginating septum. Scale bar: 200 nm. (B) Left panel, zoomed-in view of the inset in (A) in the xy coordinate plane. Right panel, same as left with PG (peptidoglycan, gray), cell membrane (peach), membrane-proximal series of dots (pink) and membrane distal series of dots (blue) highlighted. The same color scheme for labeling cellular parts is followed throughout. (C) Left panel, view of the septal disc corresponding to (B) in the xz coordinate plane obtained by rotating the cell around its short axis by 90°. Right panel, same as left with different cellular parts and cytoskeletal filaments highlighted. Scale bars for (B) and (C): 25 nm. (D) Slice through a tomogram of a dividing vegetative cell under sporulation culture conditions (see Materials and Methods). The inset highlights the left side of the leading edge of the invaginating septum. Scale bar: 200 nm. (E) Left panel, zoomed-in view of the inset in (A) in the xy coordinate plane. Right panel, same as left with different cellular parts highlighted. Densities connecting the membrane-proximal and membrane-distal series of dots are indicated in white (as pointed by a black arrow). (F) Left panel, view of the septal disc corresponding to (E) in the xz coordinate plane obtained by rotating the cell around its short axis by 90°. Right panel, same as left with different cellular parts and FtsAZ filaments highlighted. Densities connecting the membrane-proximal ring to the membrane are highlighted in yellow (as pointed by a black arrow). Scale bars for (E) and (F): 25 nm. (G) Box-and-whisker plot depicting the distribution of intensity values for the region traced by the membrane-distal and the membrane-proximal rings normalized in the range of 0 to 1. A region with approximately similar area was masked for PG, cell membrane and cytoplasm as controls. Lower intensity values (darker pixels) correspond to high mass density in the cryo-ET data.

**Fig. S3.**
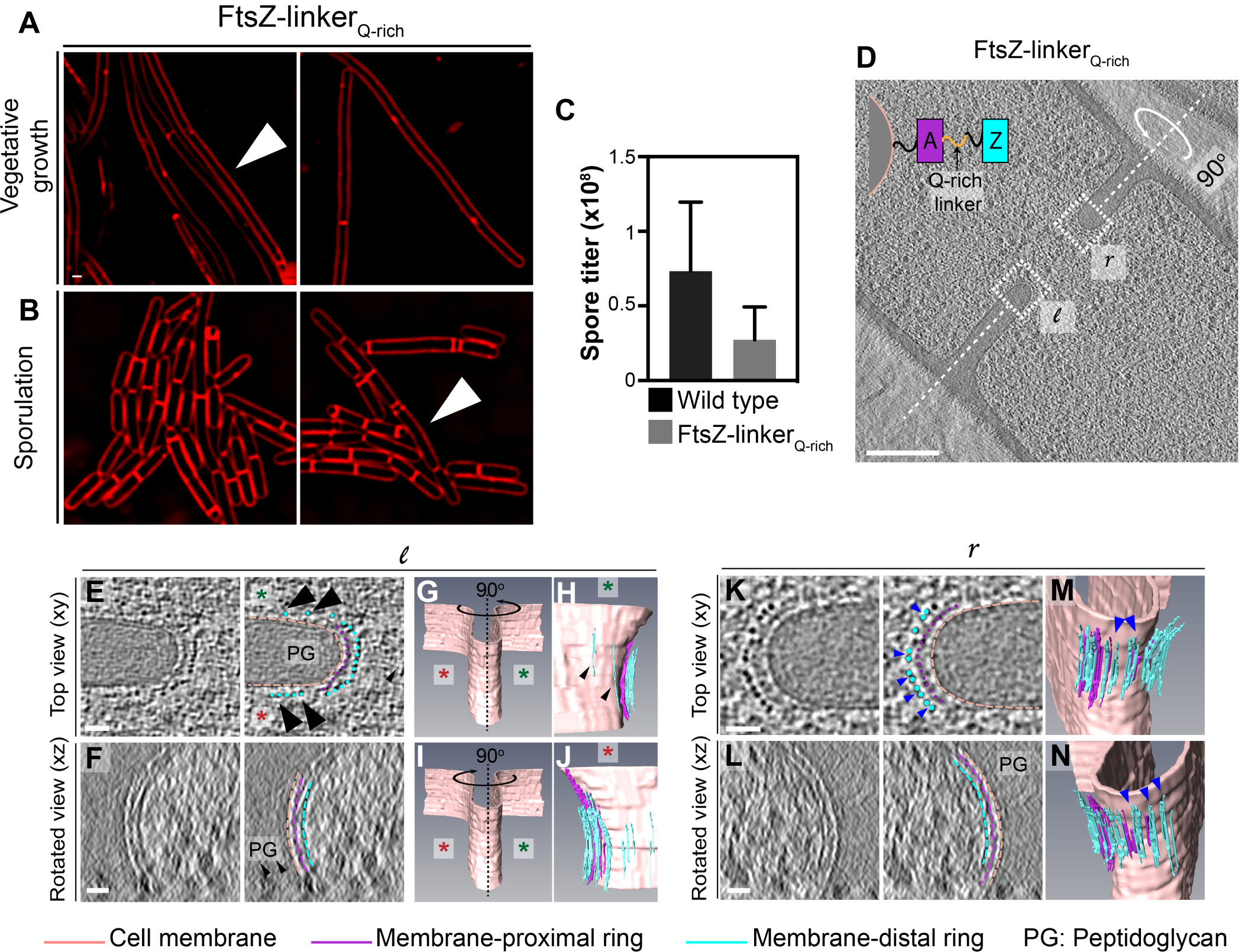
FtsZ-linker_Q-rich_ strain phenotype. (A) Morphology of FtsZ-linker_Q-rich_ strain during vegetative growth and (B) sporulation. Membranes are stained with FM4-64. The cells indicated by white arrows in (A) show a filamentous phenotype and are longer compared to the wild type. (C) Spore titer of wild type and FtsZ-linker_Q-rich_ (see Materials and Methods). (D) Slice through a tomogram of FtsZ-linker_Q-rich_ dividing cell. The insets (*l* for left and *r* for right side of the septum) highlight the leading edge of the invaginating septum. Scale bar: 200 nm. A schematic showing the construction of the modified strain is overlaid on the tomogram slice wherein PG (gray), cell membrane (peach) are highlighted. FtsZ (blue) is tethered to the membrane via FtsA (pink) and the two interact via a linker region (black + Q-rich linker in orange). The same color scheme is followed throughout the figure. (E) Left panel, zoomed-in view of the inset in (D) corresponding to ‘*l* ‘in the xy coordinate plane. Right panel, same as left with PG, cell membrane, membrane-proximal series of dots and membrane-distal series of dots highlighted. Black arrows indicate membrane-distal dots that are likely not tethered to the membrane via membrane-proximal dots. Green and red stars are used to differentiate the two opposite sides of the dividing septum. (F) Left panel, view of the septal disc corresponding to (E) in the xz coordinate plane obtained by rotating the cell around its short axis by 90°. Right panel, same as left with different cellular parts and FtsAZ filaments highlighted. Scale bar for (E) and (F): 25 nm. (G) Segmentation of the cell membrane (peach) corresponding to (E) and (F). Red and green stars indicate the two opposite sides. (H) View of the side highlighted by the green star obtained by rotating the cell by 90°as indicated in (G). Membrane-distal dots highlighted by black arrows in the right panel of (E) are highlighted. (I) Same as (G) except that the cell is rotated along its short axis by 90°to get a view of the septum side indicated by the red star. (J) View of the side highlighted by the red star. Membrane-distal dots highlighted by black arrows in the right panel of (E) are highlighted. Scale bars are omitted from (G)-(J) owing to their perspective nature. (K) Left panel, zoomed-in view of the inset in (D) corresponding to ‘*r* ‘in the xy coordinate plane. Right panel, same as left with different cellular parts highlighted. Doublets of the membrane-distal series of dots are indicated by blue arrows. (L) Left panel, view of the septal disc corresponding to (K) in the xz coordinate plane obtained by rotating the cell around its short axis by 90°. Right panel, same as left with different cellular parts and FtsAZ filaments highlighted. Scale bar for (K) and (L): 25 nm. (M) and (N) Two views of the annotated cell membrane, membrane-proximal and membrane-distal filaments corresponding to (K) and (L). Doublets of membrane-distal filaments highlighted by blue arrows in the right panel of (K) are indicated in (M) and (N).

**Fig. S4.**
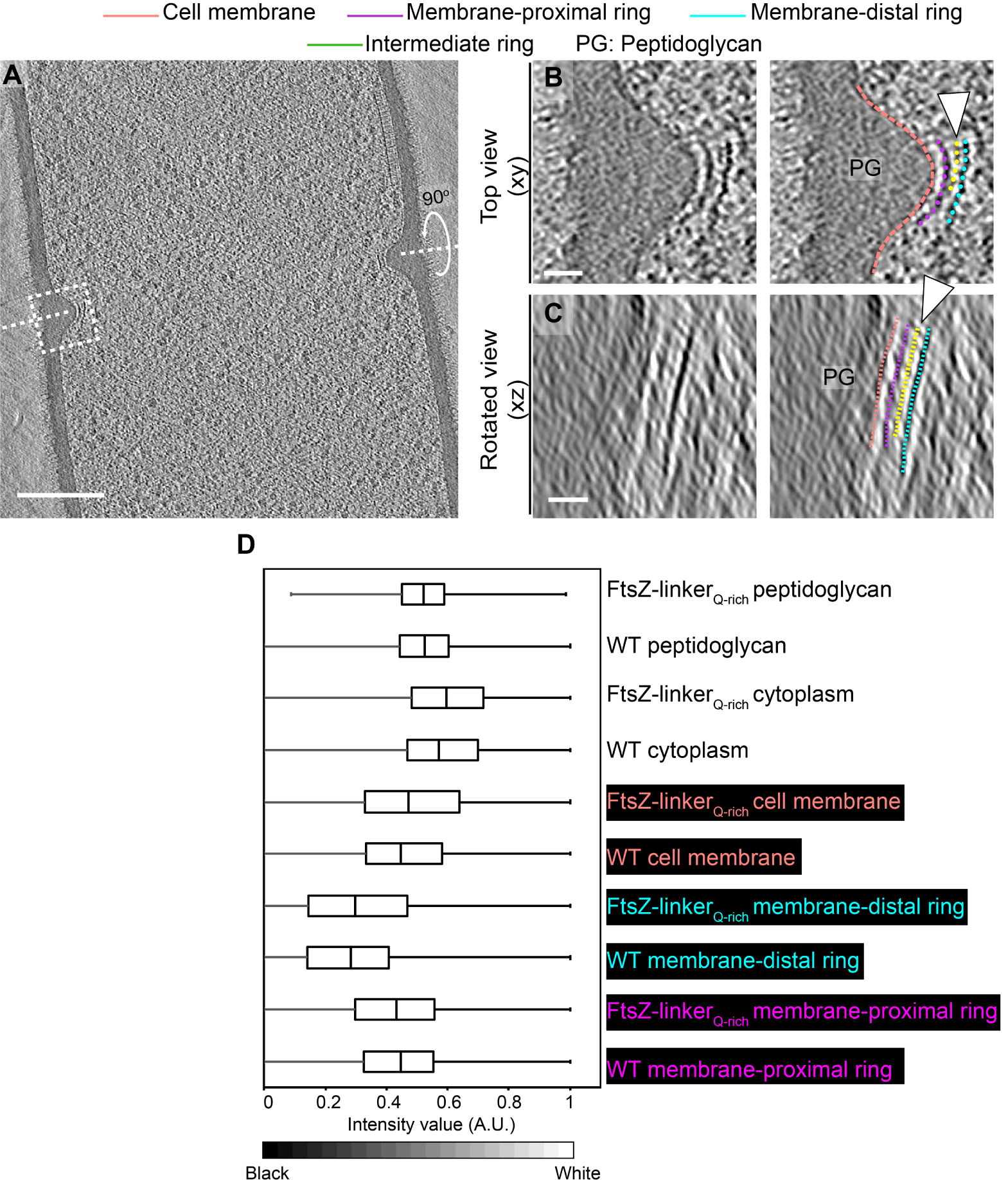
Cytoskeletal filaments in FtsZ-linker_Q-rich_. A) Slice through a tomogram of a dividing FtsZ-linker_Q-rich_ vegetative cell. The inset highlights the left side of the leading edge of the invaginating septum. Scale bar: 200 nm. (B) Left panel, zoomed-in view of the inset in (A) in the xy coordinate plane. Right panel, same as left with PG (peptidoglycan, gray), cell membrane (peach), membrane-proximal series of dots (pink) and membrane-distal series of dots (blue) highlighted. An intermediate series of dots between the membrane-proximal and the membrane-distal series is highlighted in yellow (C) Left panel, view of the septal disc corresponding to (B) in the xz coordinate plane obtained by rotating the cell around its short axis by 90°. Right panel, same as left with PG, cell membrane, membrane-proximal ring, membrane-distal ring and the intermediate ring highlighted. Scale bars for (B) and (C): 25 nm. (D) Box-and-whisker plot depicting the distribution of intensity values for the region traced by the membrane-distal and the membrane-proximal rings for wild type and FtsZ-linker_Q-rich_ normalized in the range of 0 to 1. A region with approximately similar area was masked for PG, cell membrane and cytoplasm as controls for both strains. Lower intensity values (darker pixels) correspond to high mass density in the cryo-ET data.

**Fig. S5.**
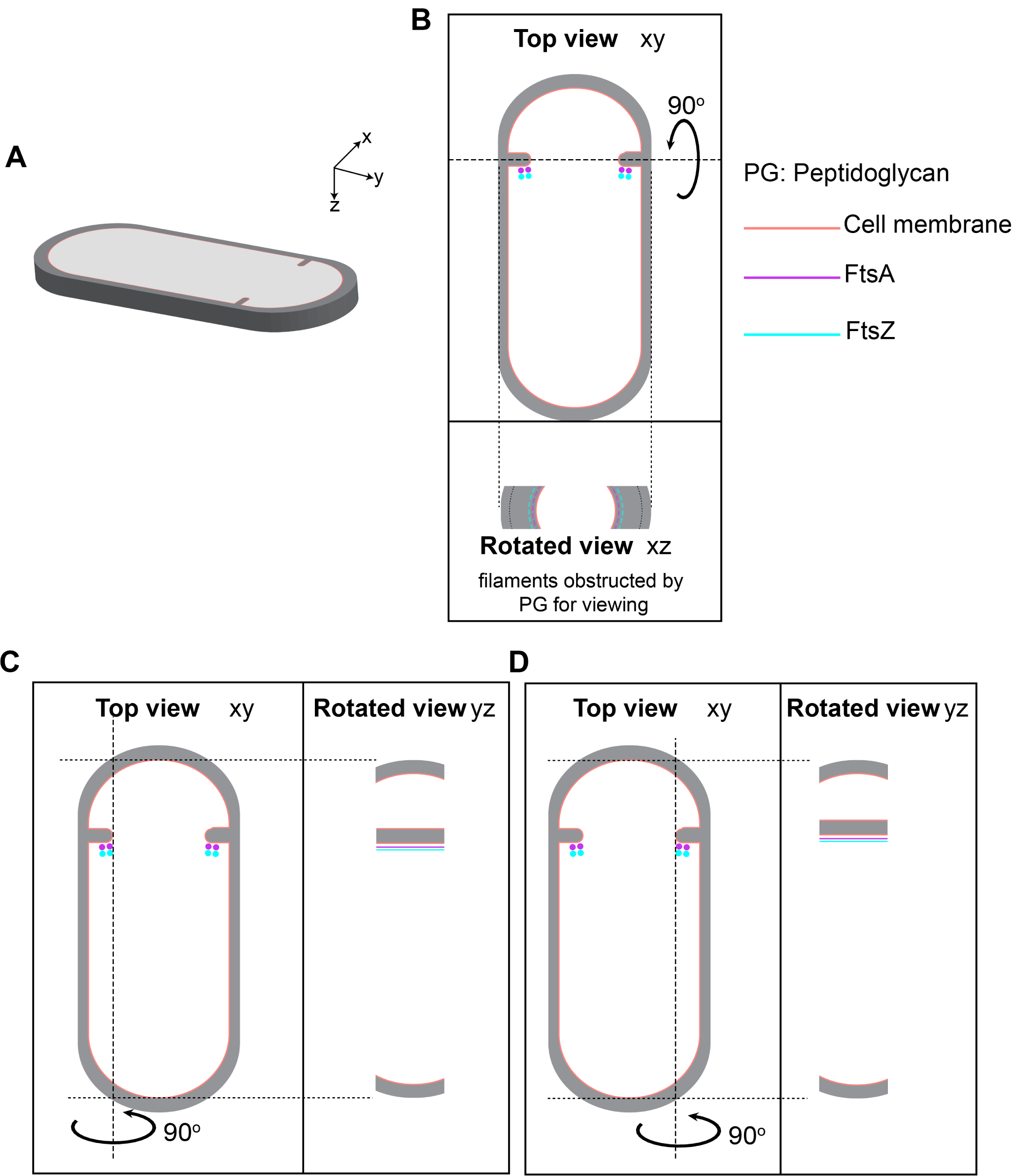
Visualizing cytoskeletal filaments in cross-sectional views in sporulating cells. (A) A section of a dividing sporulating cell. PG (grey), cell membrane (peach) are highlighted. The xyz coordinate axis to represent the dimensions of the 3D specimen is indicated (same as Fig. 1B). (B) Top panel, projection image of the cell in (A) in the xy coordinate plane. FtsA (pink) and FtsZ (blue) bundle are indicated on the mother cell side of the septum. Bottom panel, the corresponding projection image in the xz coordinate plane when the cell is rotated about its short axis by 90°. The lateral PG, septal PG and FtsAZ filaments as they project onto the orthogonal view (xz) are also highlighted. Same color scheme is followed throughout. (C) Left panel, same as top panel in (B). Right panel, the corresponding projection image in the yz coordinate plane when the cell is rotated about its long axis on the left side of the invaginating septum. (D) Left panel, same as top panel in (B). Right panel, the corresponding projection image in the yz coordinate plane when the cell is rotated about its long axis on the right side of the invaginating septum. For both (C) and (D), FtsAZ bundle and the corresponding filaments are also indicated.

**Fig. S6.**
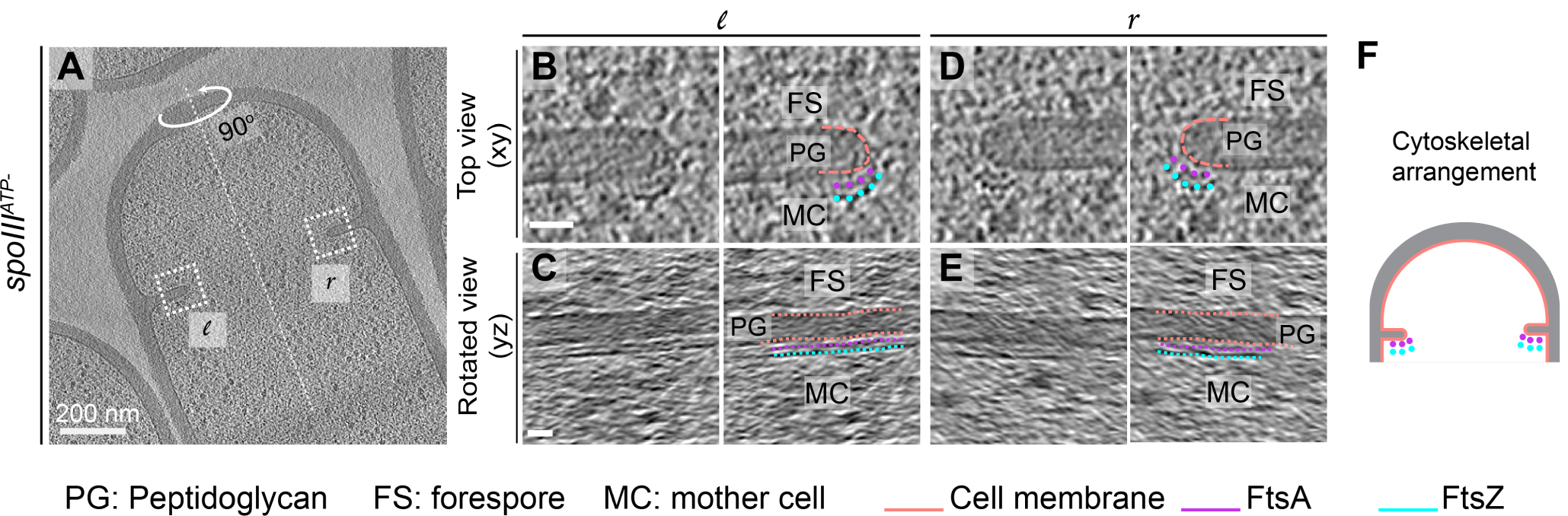
Cytoskeletal filaments in SpoIIIE^ATP-^ mutant sporangia. Somehow, we were able to better resolve FtsAZ filaments in dividing cells of SpoIIIE^**ATP-**^ - mutant sporangia during sporulation, where we observed them on the mother cell side of the septum (similar to wild type). We do not fully comprehend the reason behind this, although previous work has shown that this transmembrane protein localizes to the leading edge of the constricting membrane (43). (A) Slice through a tomogram of dividing SpoIIIEATP- mutant sporangia. The insets correspond to the left (*l*) and right (*r*) side of the invaginating septum. Scale bar: 200 nm. (B) Left panel, zoomed-in view of the inset in (A) corresponding to ‘*l*’ in the xy coordinate plane. Right panel, same as left with PG (peptidoglycan, grey), cell membrane (peach), FtsA bundle (pink) and FtsZ bundle (blue) highlighted. Same color scheme for labeling is followed throughout. (C) Left panel, view of the septal disc corresponding to (B) in the yz coordinate plane obtained by rotating the cell around its long axis near the left side of the invaginating septum by 90°. Right panel, same as left with different cellular parts and FtsAZ filaments highlighted. (D) Left panel, zoomed-in view of the inset in (A) corresponding to ‘*r* ‘in the xy coordinate plane. Right panel, same as left with different cellular parts highlighted. (E) Left panel, view of the septal disc corresponding to (D) in the yz coordinate plane obtained by rotating the cell around its long axis near the right side of the invaginating septum by 90°. Right panel, same as left with different cellular parts and FtsAZ filaments highlighted. Scale bars for (B)-(E): 25 nm. (F) Schematic showing the arrangement of the cytoskeletal machinery in dividing SpoIIIE^**ATP-**^ - sporangia with FtsAZ bundles indicated.

**Fig. S7.**
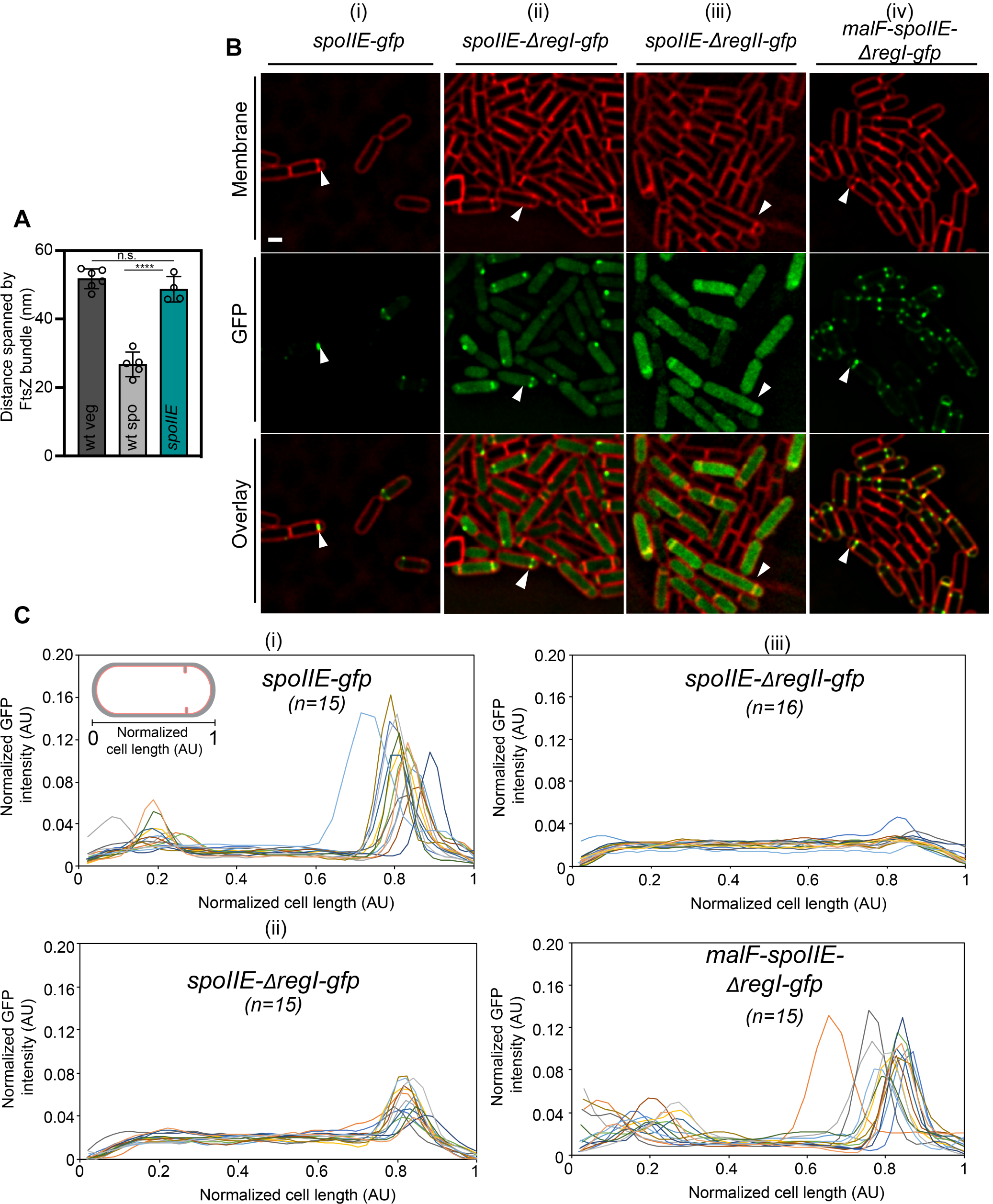
Characterization of SpoIIE mutant sporangia. (A) Bar graph depicting the distance spanned by FtsZ bundle in wild type vegetative and sporulating cells and SpoIIE null mutant sporangia. Error bars indicate standard deviation (n.s.: not significant, ****: p *≤*0.0001, unpaired t-test). Each dot indicates a sample point. (B) Fluorescence microscopy of GFP-tagged (green) (i) *spoIIE*, (ii) *spoIIE-*Δ*regI*, (iii) *spoIIE-*Δ*regII*, and (iv) *malF-spoIIE* sporangia. Membranes are stained with FM4-64 (red). Scale bar: 1 *µ*m. White arrows indicate a representative sporulating cell for each case. (C) Line graphs showing normalized GFP intensity for (i-iv) in (B) along the normalized length of sporangia. n indicates the number of cells analyzed for each case. Each line represents a single cell. In agreement with a previous study using the same strains (19), GFP signal corresponding to SpoIIE-ΔregI appeared mostly diffuse with some cells showing enrichment or punctate patterns at the polar septum (at∼ 50% level compared to wild type) indicating some association with FtsZ, GFP signal corresponding to SpoIIE-ΔregII appeared almost completely diffuse in the cytoplasm, and GFP signal corresponding to MalF-SpoIIE localized similar to wild type, displaying punctate patterns at the polar septum.

## Supplemental Movie Legends

**Movie S1**. Series of slices through the cryo-electron tomogram of a dividing vegetative *B. subtilis* cell shown in Fig. 1C. Insets highlight the leading edges of the dividing septum with cell division proteins visible as a series of dots coating the leading edge. See also Fig. 1.

**Movie S2**. Series of slices through the cryo-electron tomogram of a dividing vegetative *B. subtilis* cell shown in Fig. 1C when the cell is rotated about its short axis by 90°. The membrane-proximal and the membrane-distal rings shown in Fig. 1F,G are indicated.

**Movie S3**. Series of slices through the cryo-electron tomogram of a dividing vegetative *B. subtilis cell* shown in Fig. 1C wherein the cell membrane (peach), FtsA ring (pink) and FtsZ ring (blue) are annotated as in Fig. 1I-K.

